# Deceleration of cell cycle underpins a switch from proliferative- to terminal division in plant stomatal lineage

**DOI:** 10.1101/2021.05.17.442671

**Authors:** Soon-Ki Han, Jiyuan Yang, Machiko Arakawa, Rie Iwasaki, Tomoaki Sakamoto, Seisuke Kimura, Eun-Deok Kim, Keiko U. Torii

## Abstract

Differentiation of specialized cell types from self-renewing progenitors requires precise cell cycle control. Plant stomata are generated through asymmetric divisions of a stem-cell-like precursor meristemoid followed by the single symmetric division that creates an adjustable pore surrounded by paired guard cells. The stomatal-lineage-specific transcription factor MUTE terminates the asymmetric divisions and triggers differentiation. However, the role of cell cycle machinery in this transition remains unknown. Through time-lapse imaging, we discover that the symmetric division is slower than the asymmetric division. We identify a plant-specific cyclin-dependent kinase inhibitor, SIAMESE-RELATED4 (SMR4), as a molecular brake that decelerates cell cycle during this transition. *SMR4* is directly induced by MUTE and transiently accumulates in differentiating meristemoids. SMR4 physically and functionally associates with CYCD3;1 and extends G1-phase of asymmetric divisions. By contrast, SMR4 fails to interact with CYCD5;1, a MUTE-induced G1 cyclin, and permits the symmetric division. Our work unravels a molecular framework of the proliferation-to-differentiation switch within the stomatal lineage and suggests that a timely proliferative cell cycle is critical for the stomatal fate specification.

## INTRODUCTION

Growth and development of multicellular organisms rely on faithful cell-cycle progression, in which fundamental mechanism is highly conserved across the eukaryote kingdoms (Elledge, 1996; Harashima et al., 2013). Accumulating evidence in metazoans emphasizes that cell-cycle machinery is modulated during development, operating distinctly in proliferating stem cells vs. differentiating cells (Budirahardja and Gonczy, 2009). For example, early embryogenesis of flies, fish and frogs, as well as murine embryonic stem cells, execute rapid cell cycle mode due to shortened gap phases. As they undergo fate specification or differentiation, the duration of cell cycle increases (Coronado et al., 2013; Dalton, 2015; Liu et al., 2019). This phenomenon raises a major question regarding their causal relationships; whether the mode of cell cycle plays a role in maintaining pluripotency and whether changes in cell cycle upon dissolution of pluripotency drive fate specification or differentiation.

A typical eukaryotic cell cycle is composed of four distinct phases, G1-S-G2-M. Cell-cycle progression is driven by the oscillation of cyclin-dependent kinase (CDK) activity triggered by phase-specific cyclins, which are tightly regulated by the level of synthesis and proteolysis (Harashima et al., 2013; Morgan, 2007). CDK activity is negatively regulated by cyclin-dependent kinase inhibitors (CKIs). The G1/S transition is initiated by D-type Cyclin (CyclinD) and CDK complex, which relieve Retinoblastoma (Rb)-mediated repression on S-phase gene expression (Bertoli et al., 2013). Accumulating studies suggest that the G1 extension is indicative of differentiation (Coronado et al., 2013; Liu et al., 2019).

Plants possess a large number of genes encoding cyclins, CDKs, and CKIs (Inze and De Veylder, 2006). Studies have shown how specific cell cycle components are coupled to developmental patterning. For example, during Arabidopsis root development, transcription factors SHORTROOT and SCARECROW directly induce a D-type cyclin, CYCD6;1, that drives a formative cell division to create root endodermis and cortex cells (Sozzani et al., 2010). Another example is lateral root formation, in which auxin-induced formative division is modulated by CYCD2;1 and plant CKI, KIP-RELATED PROTEIN2 (KRP2, also known as ICK2) (Sanz et al., 2011). Some highly-specialized plant cell types, such as epidermal pavement cells and trichomes, undergo endoreduplication at the onset of terminal differentiation (Inze and De Veylder, 2006). Plant-specific CKIs, including SIAMESE (SIM) and SIAMESE-RELATED1 (SMR1, also known as LGO) regulate morphogenesis of trichomes and sepal giant cells, respectively, by promoting endoreduplication presumably via inhibiting CDK activity (Hamdoun et al., 2016; Roeder et al., 2010; Walker et al., 2000). It remains unclear if the modulation of cell cycle contributes to switching the cell division mode - from stem cell divisions to differentiating cell divisions - in plants.

Development of stomata, valves on the plant aerial epidermis for gas exchange and water control, is an accessible model of *de novo* initiation and differentiation of lineage-specific stem cells. In Arabidopsis, birth of pores begins with the stomatal-lineage fate specification of protodermal cells, which forms bipotent meristemoid mother cells (MMC) able to become either stomata or pavement cells (Han and Torii, 2016; Lau and Bergmann, 2012). A series of asymmetric cell division (ACD) follows to amplify the number of stomatal lineage precursor cells, meristemoids and stomatal lineage ground cells (SLGC). The meristemoid renews itself after ACD, thus behaving as a transient stem cell. After a few rounds of ACDs, a single round of terminal symmetric cell division (SCD) of a guard mother cell (GMC) proceeds, completing a stoma composed of paired guard cells (Han and Torii, 2016; Lau and Bergmann, 2012) (Figure 1A).

**Figure 1.**
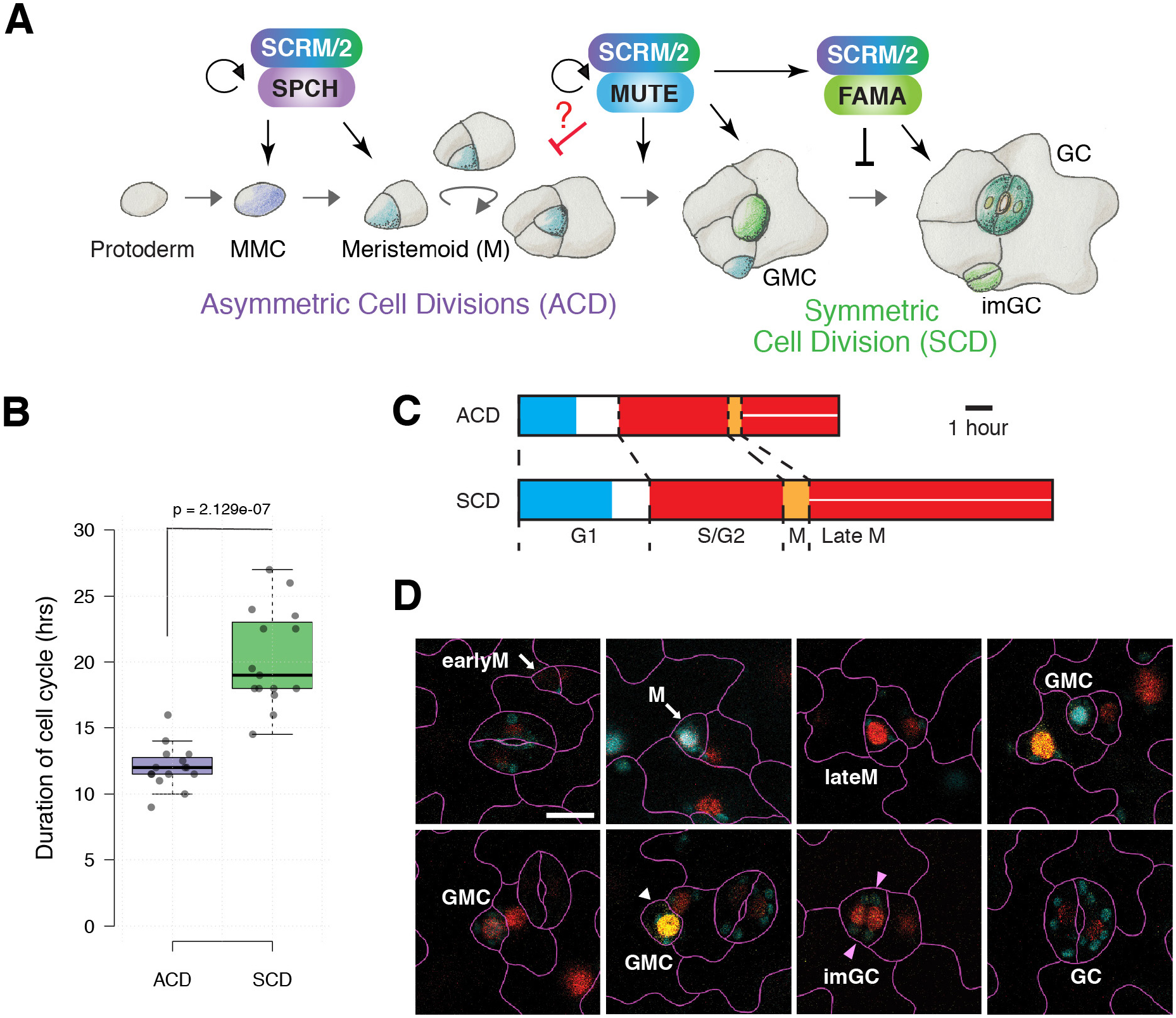
Cell cycle duration between asymmetric cell division and symmetric cell division during stomatal development. (A) Cartoon to show the key heterodimeric transcription factors for each state of stomatal development, which is accompanied by a series of asymmetric cell division (ACD) triggered by SPCH·SCRM/2 and single round of symmetric cell division (SCD) coordinated by MUTE·SCRM/2 and FAMA·SCRM/2. The molecular detail how the mode of cell cycle from ACD to SCD by MUTE·SCRM/2 has not been elucidated (red line and question mark in red). MMC: meristemoid mother cell, GMC: guard mother cell, GC: guard cell. (B) Duration of the cell cycle time of stomatal precursors undergoing ACD and SCD. n=15 for each cell division mode. Two-tailed Student t-test was performed. p=2.129e^-07^. (C) Duration of each cell cycle phase calculated from fluorescence intensity of each marker in PlaCC1. CDT1a-CFP signal (cyan) appears at the onset of G1 phase. HTR13-mCherry signal (red) accumulates when cells enter S phase and continues to late M phase. CYCB1;1-YFP signal (orange) increases in early M phase and later two separated red nuclei indicate the post division events (red with a line). Arrow: MMC; Arrowhead: GMC. Scale bar: 1 hour (D) Confocal images of stomatal lineage cells from 4-day-old cotyledon of Col-0 expressing three PlaCCI markers. M: Meristemoid; GMC: Guard mother cell; imGC: immature GC; GC: Guard cells. Scale bar: 10 μm

Accumulating evidence supports that master-regulatory bHLH proteins SPEECHLESS (SPCH), MUTE, and FAMA govern cell-state transitions within the stomatal lineage, in part via directly regulating the expression of cell cycle genes (Adrian et al., 2015; Hachez et al., 2011; Han et al., 2018; Lau et al., 2014) (Figure 1 A). SPCH initiates and sustains the ACDs of a meristemoid in part via upregulating D-type cyclins CYCD3;1 and 3;2 (MacAlister et al., 2007; Vaten et al., 2018). MUTE terminates proliferative cell state and drives final SCD by activating a large subset of cell cycle regulators, including CYCD5;1 (Han et al., 2018; Pillitteri et al., 2007). FAMA and a Myb protein, FOUR LIPS, are directly induced by MUTE and inhibit SCD via direct suppression of the cell cycle genes, thereby ensuring that the SCD occurs just once (Hachez et al., 2011; Han et al., 2018; Xie et al., 2010). However, it is not known how proliferative ACD switches to terminal SCD, and whether the core cell cycle machinery contributes to this process.

Through time-lapse imaging of stomatal development using plant cell cycle marker PlaCCI (Desvoyes et al., 2020), we discovered that the stomatal SCD cycle is slower than that of ACDs. Subsequent transcriptomic and ChIP-sequencing analyses identified that MUTE directly induces the expression of *SMR4* during meristemoid-to-GMC transition. Through loss-of-function and stomatal-lineage specific overexpression of SMR4 as well as its functional interaction studies with D-type cyclins, we elucidate that SMR4 acts as a molecular brake to decelerate cell cycle in G1 phase to ensure termination of the ACD cycle and facilitate faithful progression to SCD. This SMR4 activity is distinct from canonical functions of SIM/SMR to trigger endoreduplication. Taken together, we reveal a molecular framework of the cell proliferation-to-differentiation switch within the stomatal lineage and suggest that a timely proliferative ACD cycle is critical for differentiation of stomata with proper GC size, shape, and identity.

## RESULTS

### The single symmetric division of stomatal precursor is slower than amplifying asymmetric division

The stomatal precursor cells execute a unique transition from amplifying asymmetric cell division (ACD) to a single symmetric cell division (SCD), a step coordinated by the bHLH protein MUTE (Figure 1A). To understand if a switch from the ACD-to-SCD division mode links to the cell cycle dynamics, we first performed time-lapse imaging of developing cotyledon epidermis by using the multi-color plant cell cycle marker, PlaCCI (Desvoyes et al., 2020) and examined each phases of cell cycle during asymmetric and symmetric divisions (Figure 1B-D; Figure S1). The average cell cycle time of ACD of meristemoid and SCD of GMC was 8.63 ± 2.52 and 15.27 ± 4.73 hrs, respectively (Figure 1B), indicating that ACD is faster by ∼7.5 hours than SCD that creates a pair of guard cells. Measuring cell division time using a plasma-membrane marker LTI6b-GFP yielded essentially the same results (Figure S2). Duration of each phase was extended in the SCD (Figure 1C). On the basis these findings, we conclude that the switching from ACD to SCD involves cell cycle slow down.

### *SMR4* is expressed in stomatal-lineage cells and directly induced by MUTE

In Eukaryotic cells, CKIs negatively regulate cell-cycle progression. To identify a factor that plays a role in the decelerating cell cycle during the ACD-to-SCD transition, we surveyed publicly-available transcriptome data (Han et al., 2018; Lau et al., 2014) to search for *CKIs* that are induced by SPCH and MUTE. Among the 7 *KRP* and 17 *SIM* /*SMR* genes (Kumar and Larkin, 2017; Peres et al., 2007), only *SMR4* exhibits marked increase by the induced *MUTE* overexpression (*iMUTE*) (Figure 2A). On the other hand, a majority of *SMRs* and *KRPs* is either downregulated or unchanged upon *iMUTE* or *iSPCH* (Figure 2A). Subsequently, we performed time-course induction analysis. Consistent with the transcriptome data, *SMR4* expression was increased by iMUTE with similar kinetics to a known direct MUTE target, *TMM* (Han et al., 2018) (Figure 2B). In addition, iMUTE induced *SMR8* in a lesser extent (Figure S3). In contrast, none of the *CKIs* tested showed significant expression changes by iSPCH (Figure S3), indicating that, unlike MUTE, SPCH is not a potent inducer of *CKIs*.

**Figure 2.**
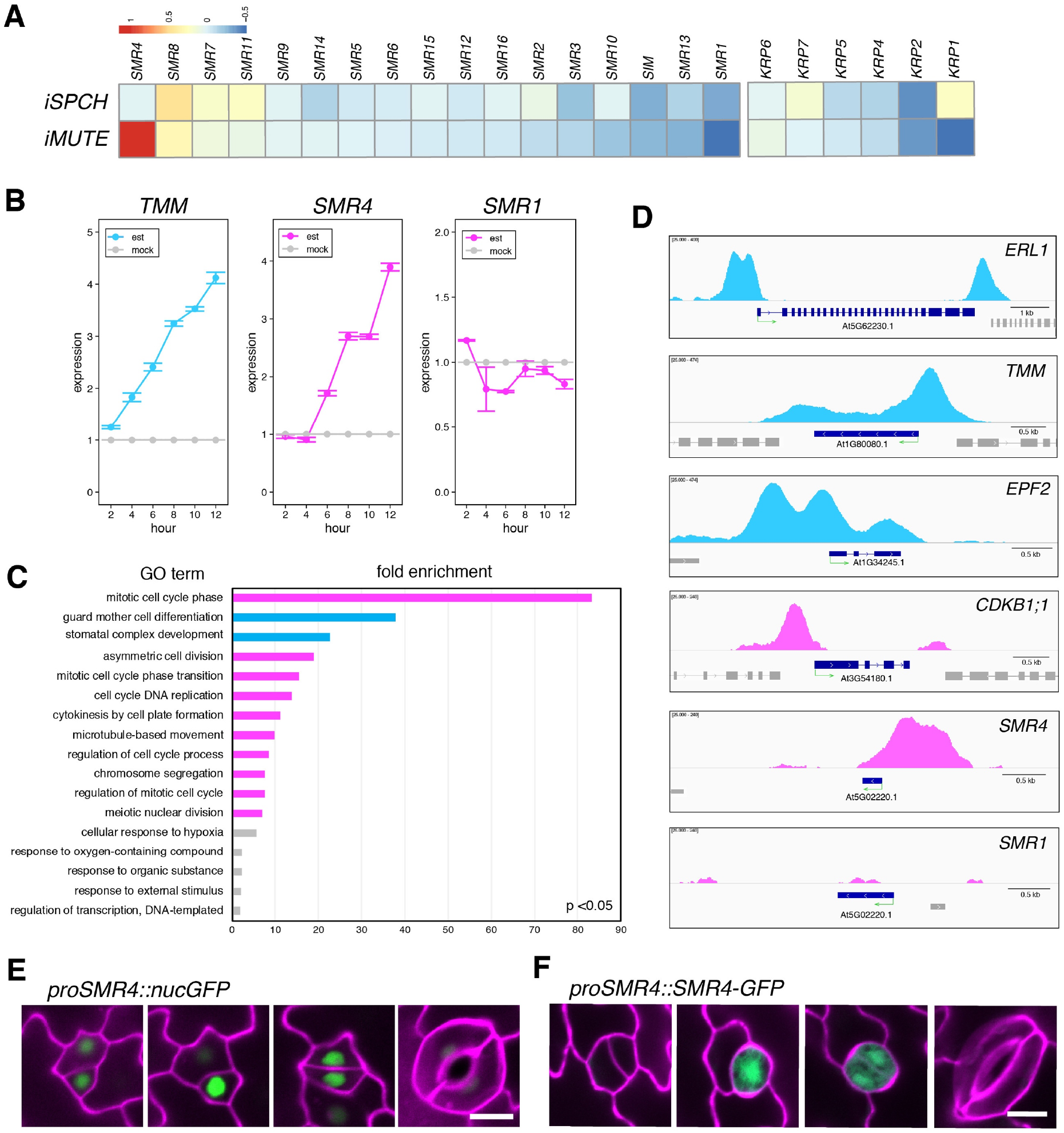
SMR4, one of the plant specific CKIs, expresses in stomatal lineage and is a direct target of MUTE. (A) Heat map represents the changes in expression of 24 CKIs in Arabidopsis by SPCH or MUTE induction. RNA-seq data adopted from Lau et al., 2014 (Lau et al., 2014) (*iSPCH*) and Han et al., 2018 (Han et al., 2018) (*iMUTE*). Heat map denotes log_2_ ratio of changes in expression compared to non-induced control. (B) Time course expression for 12 hours with 2-hour-interval of SMRs by iMUTE monitored by qRT-PCR. TMM used as a positive control as a MUTE inducible gene. est: 10 µM estradiol treated, mock: non-treated control (DMSO only). (C) GO categories of direct MUTE targets (MUTE bound, iMUTE up) ranked by fold enrichment compared to background genome. P< 0.05. Pink bars: “cell cycle”, “division”, “mitotic” categories, Blue bars: “stomatal” categories, Grey bars: others (D) IGV snapshots of ChIP-seq profile of MUTE binding to the promoter SMR4 and known MUTE targets (ERL1, TMM, EPF2, CDKB1;1) (Han et al., 2018; Qi et al., 2017). no MUTE binding was detected to *SMR1* loci. Green arrow under the gene annotation indicates gene orientation and transcriptional start sites. (E, F) Expression patterns of *SMR4* transcriptional and translational reporters. *proSMR4::nls-3xGFP* (E) and *proSMR4::SMR4-GFP* (F) in stomatal lineage precursor cell specific on the epidermis. Scale bar, 20 µm

Our previous transcriptome study (Han et al., 2018) found that MUTE induces a suite of cell-cycle- and mitotic division-related genes driving the SCD of stomata. To test whether these genes are indeed direct MUTE targets, we performed genome-wide MUTE ChIP-sequencing (see Methods) (Table S1). The overwhelming majority of MUTE-bound genes (Table S2) that are upregulated by MUTE belongs to the Gene Ontology (GO) categories (Table S3).: “mitotic cell cycle phase (83.22-fold enrichment, p=2.15e-02)”, “asymmetric cell division (18.91-fold enrichment, p=2.4-e-02)” and other cell cycle/mitosis related categories (Figure 2C, pink bars), as well as the genes involved in stomatal development: “guard mother cell differentiation (37.83-fold enrichment, p=8.34e-04)” and “stomatal complex development (22.7-fold enrichment, p=1.75e-12)” (Figure 2C, cyan bars). Strong MUTE-bound peaks are detected at the 5’- and 3’-regions of known MUTE target loci, *ERL1, TMM, EPF2*, and *CDKB1;1* (Figure 2D). Most importantly, MUTE robustly bound to the 5’ region of *SMR4*, indicating that *SMR4* is a direct MUTE target. As expected, no MUTE binding peak was detected in *SMR1* loci, which is not induced by iMUTE and thus not a direct MUTE target (Figure 2B, D).

We further characterized the *SMR4* expression patterns using seedlings expressing nuclear-localized GFP driven by the *SMR4* promoter (Figure 2E). A strong GFP signal was detected in the stomatal-lineage cells with the highest expression in a late meristemoid transitioning to GMC, and persisted after SCD (Figure 2E). Likewise, a translational reporter of SMR4-GFP fusion protein driven by the *SMR4* promoter exhibited similar accumulation patterns predominantly in the nuclei (Figure 2F). These expression patterns mirror that of MUTE (Pillitteri et al., 2007) Combined, our results indicate that MUTE directly promotes the *SMR4* expression in stomatal precursor cells before the onset of the SCD. Furthermore, the *SMR4* expression suggests its distinct role from canonical CKIs.

### *SMR4* suppresses cell proliferation in part with *SMR8*

To understand the role of *SMR4* in stomatal development, we next sought to characterize its loss-of-function phenotypes. Because no T-DNA insertion line is available for *SMR4*, presumably owing to its short coding sequence (219 bp), we employed CRISPR-Cas9 system (Tsutsui and Higashiyama, 2017)(see Methods). A guide RNA targeting to *SMR4* yielded either a base-pair deletion (*smr4-1cr*) or insertion (*smr4-2cr*) at 80 bp from the translation start site, which leads to a frame shift and premature stop codon (Figure S4A, B). A quantitative analysis of segmented epidermal cells (see Methods) revealed that *smr4cr* epidermis are increased in small cells (< 50 μm^2^) and concomitantly decreased in large pavement cells (> 4000 μm^2^) (Figure 3A, B). Stomatal precursor cell density is also increased in *smr4cr* alleles (Figure 3C). Introduction of functional *SMR4* transgene (*proSMR4::HA-SMR4*) fully rescued the phenotypes of *smr4-1cr* (Figure S4D-G), indicating that increased numbers of stomata and stomatal precursor cells in *smr4* are due to the loss of function of *SMR4*.

**Figure 3.**
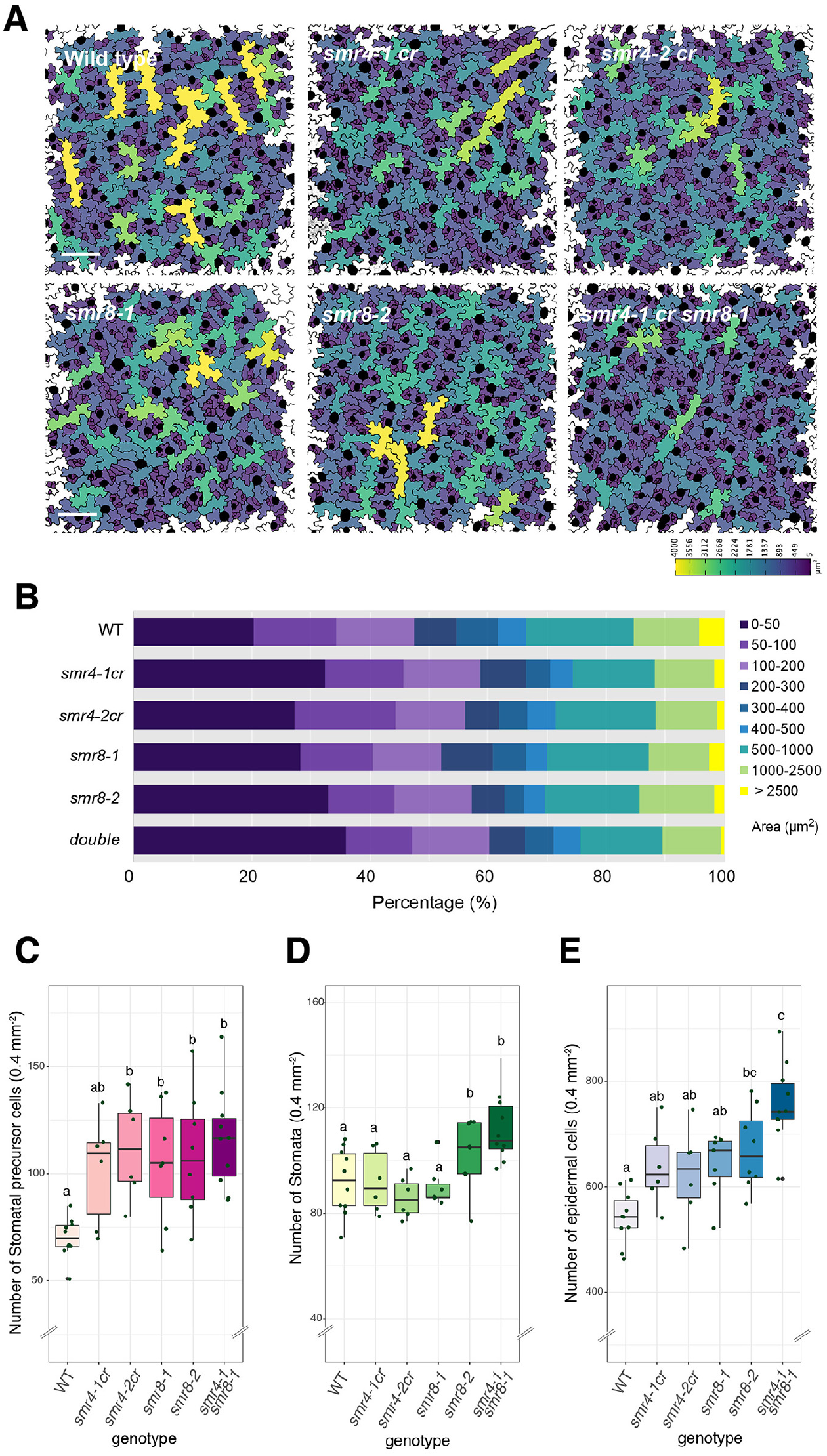
*smr4* CRISPR knockout mutants produce smaller cells, the phenotype enhanced by *smr8*. (A) Abaxial epidermis of cotyledons at 4-day-old seedlings of wild-type, *smr4-1cr, smr4-2*cr *smr8-1* and *smr4-1 smr8-1* double mutant Arabidopsis plants. Epidermal cells size is color-coded as a color scale at bottom. Scale bar, 100 μm (B) Bar graphs showing the percentage of each category of cell area from the images for the genotype presented in (A). (C-E) Quantification data of the number of stomatal precursor cells (meristemoid + GMC) (C), stomata (D), and total epidermal cells (E) per 0.4mm^-2^ area for the genotypes shown in (A). One-way ANOVA followed by Tukey’s post hoc test was performed for comparing all genotypes. Different letter denotes significant difference. Double letter denotes insignificance. P <0.05 or P<0.01. The number of plants from each genotype, WT: n=10, *smr4-1cr*: n=6, *smr4-2cr*: n=6, *smr8-1*: n=7, *smr8-2*: n= 8, *smr8-1 smr8-2*: n=10

Because *SMR8* expression was marginally increased by iMUTE (Figure S3), we further characterized the loss of function phenotypes of *SMR8*. Two T-DNA insertion lines, *smr8-1* and *smr8-2*, accumulate a reduced-level of *SMR8* transcripts (Figure S4C). Like *smr4, smr8* mutants conferred an increase in small epidermal and stomatal precursor cells, and this phenotype was most exaggerated in the *smr4 smr8* double mutant (Figure 3A, B, D, E). Therefore, *SMR4* plays a role in repressing ACD, including amplifying- and spacing cell division of precursor cells, in part redundantly with SMR8.

### Stomatal-lineage specific expression of CKIs reveals their unique functions

Our study revealed that *SMR4* is a direct MUTE target and expresses during the transition from proliferating meristemoid state to differentiating GMC state. SMR proteins are known to promote endoreduplication in trichomes, pavement cells and sepal giant cells (Hamdoun et al., 2016; Kumar and Larkin, 2017; Roeder et al., 2010; Walker et al., 2000). However, unlike trichomes and pavement cells, stomatal lineage cells do not undergo endoreduplication (Melaragno et al., 1993). We thus hypothesized that SMR4 may function differently from the other canonical SMRs. To address this, *SMR4, SMR8*, and *SMR1* along with *KRP1* are ectopically expressed in the stomatal lineage cells (MMC, meristemoids and SLGC) by using *POLAR* promoter (Pillitteri et al., 2011) (Figure 4; Figure S5). Unlike SMR1 or KRP1, *POLAR-*promoter-driven SMR4 and SMR8 did not significantly changed stomatal index (number of stomata /(number of stomata and non-stomatal epidermal cells) x100) (Figure 4F), reflecting reduction in the number of both stomata and epidermal cells (Figure 4G, H). Whereas GMCs of *POLAR*-promoter-driven SMR4 executed SCD, they formed stomata composed of skewed guard cells (Figure 4B, pink bracket). This suggests that SMR4 does not inhibit the final SCD *per se*. Similar deformed stomata were also observed in *proPOLAR::SMR8* (Figure 4C, pink bracket).

**Figure 4.**
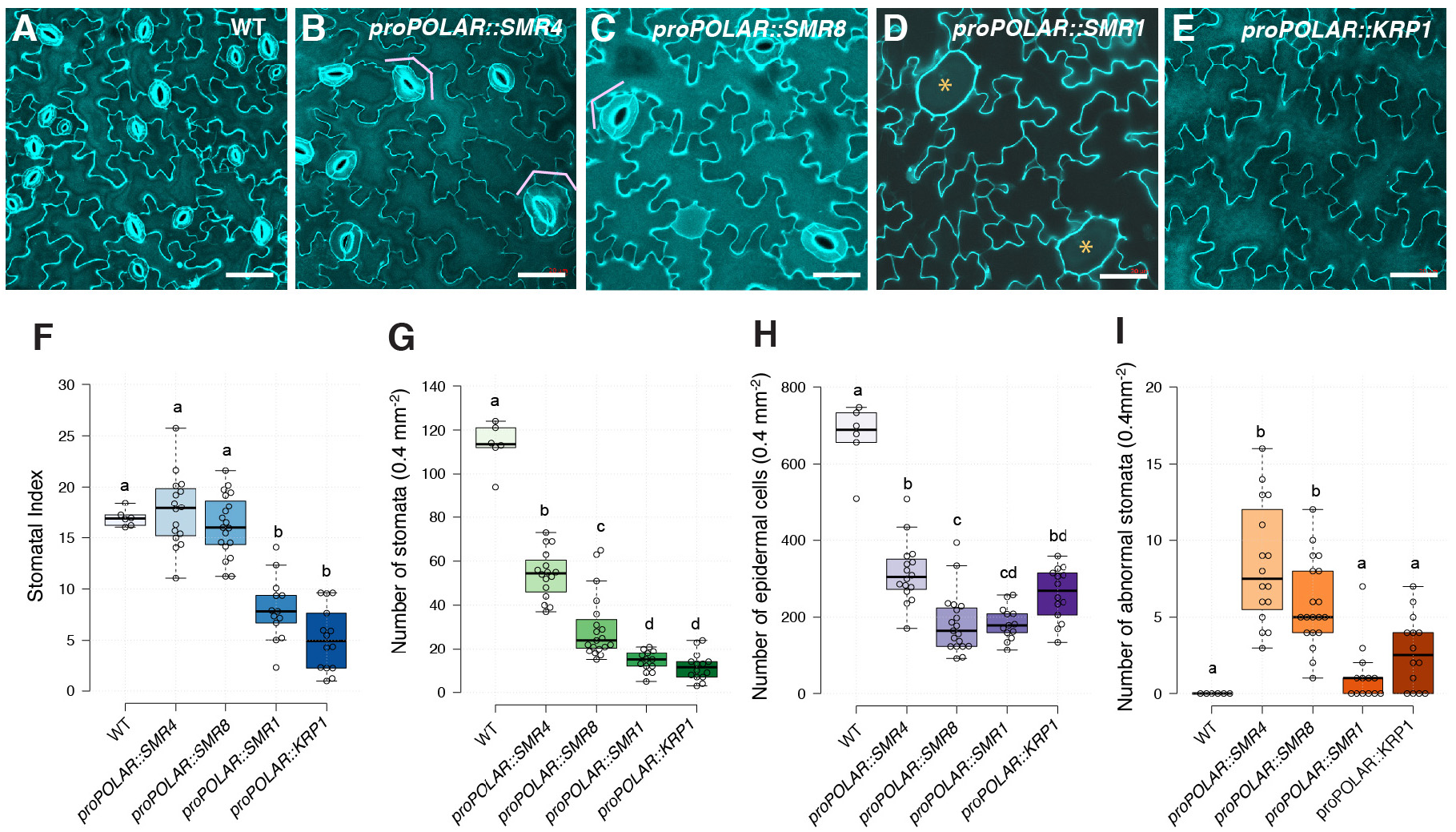
Stomatal-lineage overexpression phenotype of a suite of CKIs reveal their unique activities. (A-E) Epidermal phenotype of abaxial cotyledon in 8-day old wild type (WT: A), *proPOLAR::SMR4* (B), *proPOLAR::SMR8* (C), *proPOLAR::SMR1* (D), *proPOLAR::KRP1* (E). Scale bar, 50 µm; Orange asterisks, undivided single-celled stomata (F-I) Quantification of epidermal cell number of abaxial cotyledon in 4-day old WT and transgenic plants. Stomatal Index (F) as well as number of stomata (G) number of epidermal cells (H), and number of abnormal stomata (I) in 0.4mm^-2^ area. One-way ANOVA with Tukey’s post hoc test was performed to compare all genotypes. The number of plants from each genotype, WT n=6, *proPOLAR::SMR4* n=16, *proPOLAR::SMR8* n=19, *proPOLAR::SMR1* n=13, *proPOLAR::KRP1* n= 14

Unlike SMR4, ectopic stomatal-lineage expression of SMR1 and KRP1 displayed cell division defects with unique consequences. *proPOLAR::SMR1* produced abnormally large undivided GMC-like cells (Figure 4D, asterisks; Figure S5), consistent with the known role of SMR1 in suppressing the activity CDKB1;1 thereby promoting endoreduplication (Kumar et al., 2015). Finally, *proPOLAR::KRP1* severely inhibited the ACDs, resulting in epidermis vastly consisted of pavement cells with low stomatal index (Figure 4E, F), resembling with *spch* mutant (MacAlister et al., 2007; Pillitteri et al., 2007).

The phenotype of *proPOLAR::SMR4* (and *SMR8*) is consistent with the diminished proliferative activity of meristemoids. To delineate formative step from differentiation, we introduced *proPOLAR::SMR4* into *mute* mutant, in which a number of amplifying ACDs are increased and meristemoids arrest (Figure S5). Indeed, *proPOLAR::SMR4* significantly reduced the number of ACDs, resulted in low number of larger meristemoids (Figure S5F-J). Taken together, we conclude that SMR4, and some extent SMR8, possesses a unique feature different from canonical CKIs to specifically terminate (but not fully inhibit) ACDs of meristemoids but allows final SCD to proceed in GMC. Furthermore, formation of skewed irregular-shaped stomata, some resembling pavement cells (Figure 4B), implies that excessive SMR4 activity disrupt guard cell morphogenesis.

### SMR4 balances between cell proliferation and differentiation

To further understand the identity of abnormally-shaped stomata observed in *proPOLAR::SMR4*, stomatal lineage markers were introduced. In wild type, stomatal lineage precursor marker *proTMM::GUS-GFP* (Nadeau and Sack, 2002) was detected in stomatal lineage cells with the brightest signal in triangular shaped meristemoids (Figure 5A). Surprisingly, both stomatal-lineage cells and enlarged pavement cell-like cells in *proPOLAR::SMR4* plants expressed *proTMM::GUS-GFP* (Figure 5B). These GFP+ cells in *proPOLAR::SMR4* plants sometimes divide symmetrically (Figure 5B, E, F cyan arrows). Further quantitative analysis shows that, compared to wild type, these *proTMM::GUS-GFP* positive cells in *proPOLAR::SMR4* are greater in size range (50∼800 μm^2^) and in addition have low circularity (Figure 5C). This might reflect mixed cell fate; pavement cell shape with stomatal identity. Some of the these cells express stomatal fate commitment marker *proMUTE::nucYFP* (Figure 5F) and finally differentiate into mature guard cells (Figure 5G-I) exhibiting large wavy and skewed shape (Figure 5H, I) but expressing a mature guard cell marker, E994.

**Figure 5.**
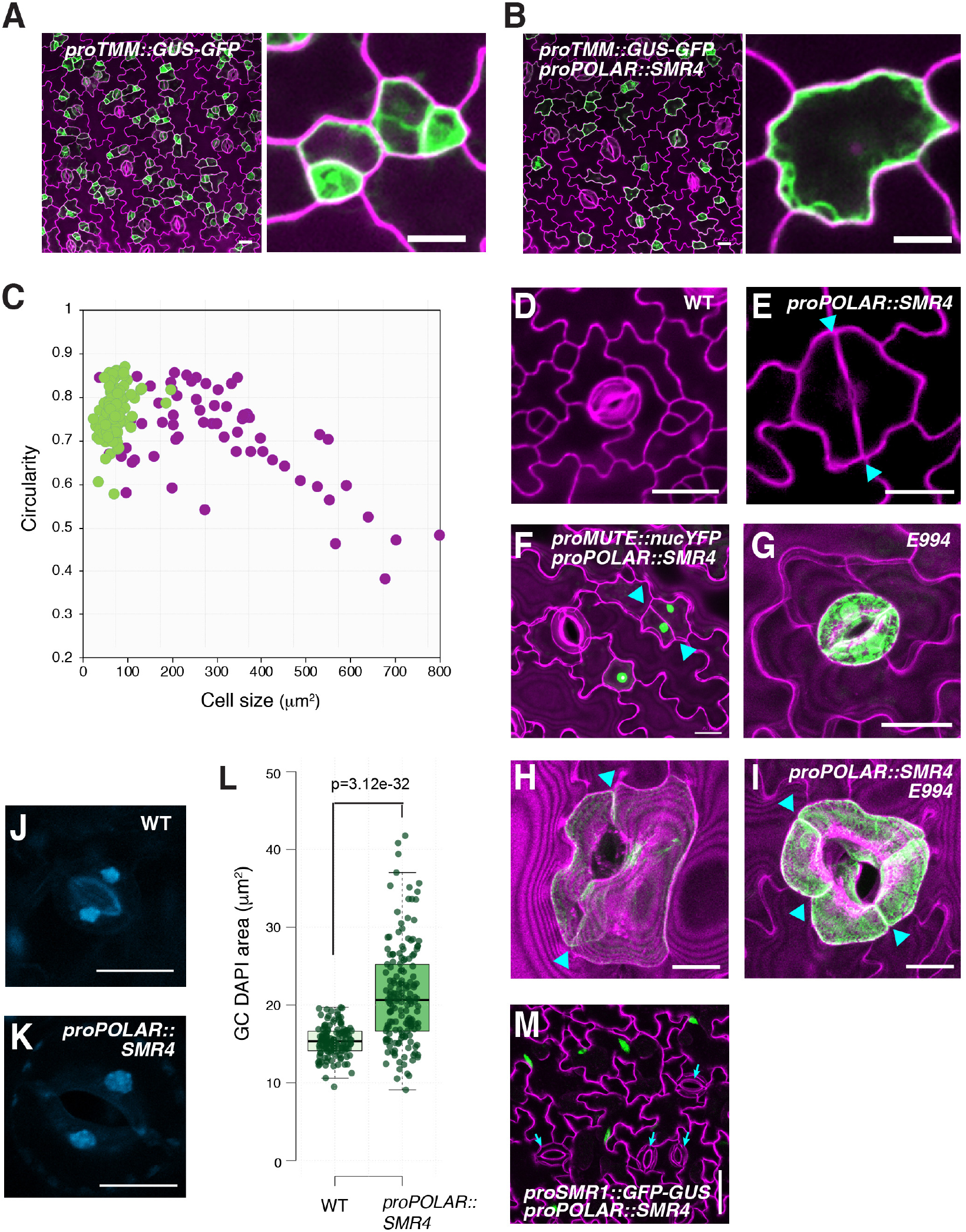
Stomatal-lineage overexpression of *SMR4* reduces proliferative activity of meristemoids. (A) *proTMM::GUS-GFP*, abaxial cotyledon at 4-day post germination stage seedling (4dpg), (B) *proTMM::GUS-GFP* in *proPOLAR::SMR4*, 4dpg, Scale bar: 20 µm. Insets: zoomed stomatal lineage cells expressing GFP. Scale bar, 10 µm (C) Size distribution vs. circularity of the stomatal lineage precursor cells expressing *proTMM::GUS-GFP* in wild-type (green dots) and *proPOLAR::SMR4* (purple dots) plants. (D-I) Confocal images of representative stomata: Wild-type stoma (D); Mixed fate stoma developed in *proPOLAR::SMR4* (E); *proMUTE::nYFP* in *proPOLAR::SMR4* (F); Mature GC marker *E994* in wild type (G) and *proPOLAR::SMR4* (H, I). Cyan arrowheads, division site of GCs. Scale bars, 20 µm (J, K) DAPI stained nuclei in mature GCs from wild-type (J) and *proPOLAR::SMR4* plants (K). (L) Quantitative analysis of DAPI-stained nuclear area in wild-type (n=128) and *proPOLAR::SMR4* (n=234) GCs. Two-tailed Student t-test was performed. p=3.12e^-32^. (M) Endoreduplication marker *proSMR1::GFP-GUS* expression in *proPOLAR::SMR4* plants. White arrows indicate enlarged GCs with no GFP expression. Scale bar, 50 µm

To address whether these enlarged GCs undergo endoreduplication process when *SMR4* is ectopically expressed, the DNA content was measured using DAPI fluorescence. Half of GC populations exhibited the fluorescence similar to the wild-type (the median value is 15 µm^2^ in wild type and 20 µm^2,^ in *proPOLAR::SMR4* plants) whereas some showed fluorescence values nearly doubled in *proPOLAR::SMR4* plants (Figure 5J-L). Likewise, the GC nuclei size measurements using H2B-GFP (Maruyama et al., 2013) are consistent with the DAPI measurements (Figure S6). The difference in nuclei size was more pronounced in pavement cells. Furthermore, none of the GCs express the endoreduplication marker *proSMR1::nlsGFP-GUS* (Bhosale et al., 2018) regardless of the GC size in *proPOLAR::SMR4* plants, whereas pavement cells, where endoreduplication normally occurs, express GFP signal (Figure 5M). Combined, these results suggest that stomatal-lineage overexprssion of *SMR4* induces another round of DNA synthesis but does not trigger the endoreduplication cycle in the stomatal lineage. This feature distinguishes SMR4 from the known SIM/SMR-family of CKIs. The unique, non-canonical activity of SMR4 is also supported by systematic stomatal-lineage overexpression of selected CKIs, where only SMR1 generated huge undivided GMC cells (Figure 4D, Figure S5).

### SMR4 decelerates cell cycle progression by G1 phase extension

We elucidated that stomatal-lineage ACDs are faster than the final SCD (Figure 1B, C). What is the ramification of stomatal-lineage overexpression of *SMR4* on cell cycle duration of ACDs and SCD? To address this question, we introduced the triple cell cycle marker PlaCCI to *proPOLAR::SMR4* plants and performed time-lapse imaging (Figure 6; Movie S3). The stomatal precursor cells (meristemoids) in *proPOLAR::SMR4* seedlings exhibited striking extension of G1 phase (Figure 6A, from average of 3.23 to 8.03 hours), extending the total cell cycle time from 12.03 to 18.47 hours (Figure 6B). In contrast, the cell cycle duration of the SCD was not significantly affected by *proPOLAR::SMR4* (19.57 hours, Figure 6, Movie S4). Combined, the results suggest that SMR4 can slow down cell cycle progress by G1 phase extension, and that the cell cycle of proliferative ACDs, but not the final SCD, are preferentially decelerated by SMR4.

**Figure 6.**
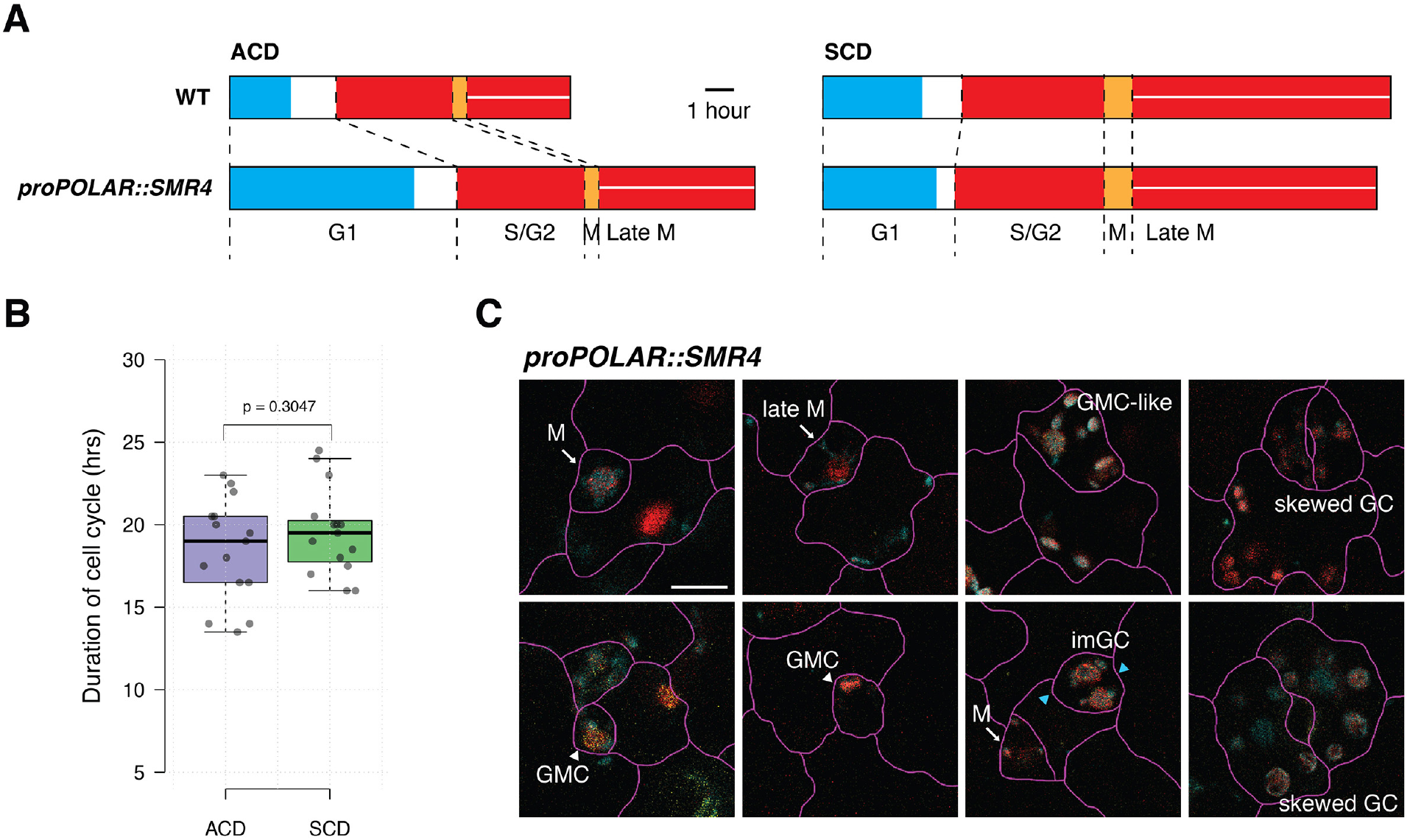
SMR4 slows down the cell cycle progression of ACD through G1 extension. (A) Duration of the cell cycle time of stomatal precursors undergoing ACD and SCD in *proPOLAR::SMR4* lines. n=15 for each cell division mode. Scale bar, 1 hour (B) Duration of each cell cycle phase calculated from fluorescence intensity of each marker. n=15 for each cell division mode. (C) Confocal images of stomatal lineage cells from 4-day old cotyledons of a plant expressing three PlaCCI markers. Colors represents the same cell cycle phase as in Fig.1C. Arrow: MMC; Arrowhead: GMC. Scale bar, 10 μm

### Physical and functional interactions of SMR4 with D-type cyclins underscore the swith from ACD to SCD

SIM is known to interact with CYCA2;3 to promote endoreduplication (Wang et al., 2020). Unlike SIM, stomatal-lineage overexpression of *SMR4* can extend G1 cycles of ACD but allows execution of the final SCD (Figures 4-6). We thus predict that SMR4 regulates the G1-S transition via associating with D-type Cyclins. To understand the mode of action of SMR4, we first surveyed publicly available protein-protein interactome data (Szklarczyk et al., 2019) (Figure 7A). The known SMR4 interactors include major components of cell cycle progression, CSK1, CKS2, CDKA;1 (CDC2), CYCD2;1, CYCD3;1 and CYCD7;1 (Figure 7A). Among them CYCD3;1 is known to play a role in SPCH-mediated ACDs, and CYCD7;1 is involved in the SCD of GMC (Adrian et al., 2015). Our yeast-two-hybrid (Y2H) analysis shows that consistent with the interactome data (Figure 7A), SMR4 associates with CYCD3;1 and CYCD7;1. In contrast, no interaction was observed for SMR4 and CYCD5;1, a direct MUTE target initiating the single SCD (Han et al., 2018) (Figure 7B). We also did not observe the direct interaction of SMR4 with CDKA;1 or CDKB1;1 (Figure 7B).

**Figure 7.**
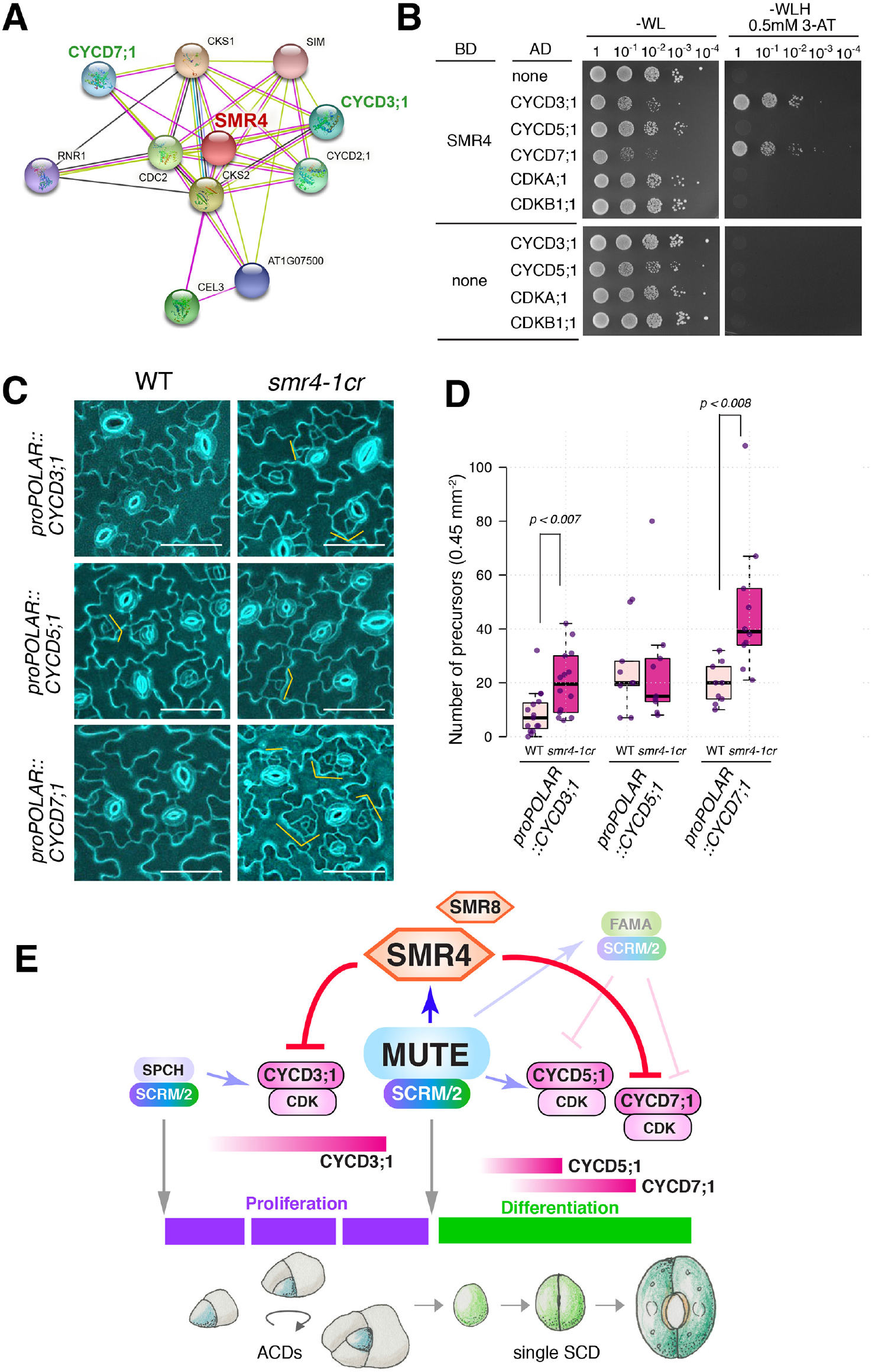
SMR4 decelerates the cell cycle via direct interactions with a selected set of D-type Cyclins. (A) SMR4 interacting proteins visualized by cytoscape. (B) Yeast-two hybrid assays in which the DNA-binding domain alone (BD) alone or fused to SMR4 were used as bait, and the activation domain alone (AD) or fused to three D-type cyclins, CYCD3;1, CYCD5;1, CYCD7;1 and two cyclin dependent kinases, CDKA;1 and CDKB1;1. Transformed yeast were spotted in 10-fold serial dilutions on appropriate selection media. (C) Transgenic plants harboring CYCD3;1, CYCD5;1 and CYCD7;1 driven by *POLAR* promoter in wild type and *smr4-1cr*. Scale bar: 50 µm (D) Quantification of stomatal precursor cells (meristemoids and GMC). Epidermal cells were counted from abaxial cotyledon of 7 dpg seedling. Mann-whitney test was performed. P-value of statistically significant pair was marked on top of the boxplot. Independent T1 transgenic plants were analyzed, the number of plants used as follows. *proPOLAR::CYCD3*;1: n=15, *proPOLAR::CYCD3*;1 *smr4-1cr*: n=14, *proPOLAR::CYCD5*;1: n=9, *proPOLAR::CYCD5*;1 *smr4-1cr*: n=9. *proPOLAR::CYCD7*;1: n=9, *proPOLAR::CYCD7*;1 *smr4-1cr*: n=10. *proPOLAR::CYCD7*;1: n=12, *proPOLAR::CYCD7*;1 *smr4-1cr*: n=12 (E) Schematic model. SPCH·SCRM/2 module initiates and sustains ACD and MUTE·SCRM/2 trigger SCD (grey arrows) by transcriptionally activating *CYCD3;1* and *CYCD5;1* (shaded blue arrows), respectively. MUTE directly up-regulates *SMR4* transcription (Blue arrow). SMR4 (and SMR8 in part) suppress the activity of CYCD3;1 and CYCD7;1 complexed with CDKs, (red line), but not CYCD5;1, to terminate ACD mode and ensure faithful progression of SCD.

Next, to address the biological significance of SMR4 interactions with CYCD3;1 and CYCD7;1 but not with CYC5;1, we examined the effects conferred by stomatal-lineage overexpression of three D-type Cyclins in the presence or absence of functional *SMR4*. As shown in Figure 7C and D, in the absence of *SMR4, POLAR*-promoter-driven expression of *CYCD3;1* and *CYCD7;1* exaggerated the ACDs, resulting in significant increase in the number of stomatal precursor cells. By contrast, *smr4-1cr* mutant did not affect the phenotype of *proPOLAR::CYCD5;1* (Figure 7C, D). On the basis of these findings, we conclude that SRM4 can suppress the stomatal-lineage divisions by direct association with CYCD3;1 and CYCD7;1, but not with CYCD5;1, and this differential interaction with D-type cyclins underscores the transition from proliferative ACDs to final SCD (Figure 7E).

## Discussion

In this study, we discovered that proliferative ACDs has faster cell cycle duration than the single terminal SCD within the stomatal cell lineages. A subsequent genome-wide profiling of MUTE targets followed by phenotypic and functional characterizations identified SMR4 as a non-canonical CKI that sets a cell-cycle break to facilitate transition from ACD to SCD. SMR4 is a direct MUTE target specifically induced by MUTE but not by SPCH (Figure 2), thus highlighting the orchestration of cell-state switch from proliferation (meristemoids) to differentiation (stomata) at the control of cell cycle duration. This view is further supported by the findings that prolonged G1/S phase specifically during the proliferative ACDs by stomatal-lineage overexpression of *SMR4* causes mis-specification of guard cells (Figures 4, 5).

In contrast of SIM and SMR1, known regulators of endoreduplication (Roeder et al., 2010; Wang et al., 2020), we found that SMR4 extends the G1/S transition during stomatal ACDs (Figure 5). It has been shown that SIM associates with CYCA2;3 but not with CYCD3;1 (Wang et al., 2020). Assuming that SMR1 functions similarly to SIM, the enlarged single-celled stomata conferred by the stomatal-lineage overexpression of SMR1 (Figure 4) can be attributed to the direct inhibition of CYCA2;3-CDKB;1 complex by SMR1. Indeed, higher-order mutations in *CYCA2s* (*cycA2;1, 2;2, 2,3* triple mutant) as well as the dominant-negative inhibition of *CDKB1;1* exhibit the identical, single-celled stomata phenotypes (Boudolf et al., 2004; Vanneste et al., 2011). By contrast, we found that SMR4 functionally associates with D-type Cyclins (Figure 7). Thus, distinct functions among SIM/SMRs lie in their unique interaction potential with different Cyclin-CDK complexes. During mammalian cell cycle, a series of CKIs exhibit inhibitory roles during G1/S transition via associating with CyclinD1/2/3-CDK4/6 and then with CyclinE/CDK2 complexes (Sherr and Roberts, 2004). Among them, p27^KIP1^ can bind with different Cyclin-CDK complexes and exert different regulatory effects on each complex (Sherr and Roberts, 2004). Plants lack CyclinE, but the previous large-scale expression analysis of cell cycle genes suggests that the plant CYCDs adopt both metazoan CyclinD and CyclinE functions (Menges et al., 2005). That SMR4 binds with different D-type Cyclins to negatively regulate G1/S phase therefore echoes its functional parallel to metazoan CKI, p27^KIP1^.

How could SMR4 decelerate cell cycles in proliferative ACDs but not in terminal SCD? Our results suggest that the specificity lies on preferential association of SMR4 with different CyclinDs, each with a unique expression pattern within the stomatal-cell lineages. For example, *CYCD3;1* and *CYCD3;2* are induced by *SPCH* and promote ACDs (Adrian et al., 2015; Han et al., 2018; Lau et al., 2014). By contrast, *CYCD5;1* is directly induced by MUTE to drive the terminal SCD (Han et al., 2018). *CYCD7;1* is likely involved in the terminal SCD, however, its expression starts later and persists longer than *CYCD5;1* (Han et al., 2018; Weimer et al., 2018). Based on the physical and functional associations of SMR4 with CYCD3;1 and CYCD7;1 but not with CYCD5;1, we propose the following model of regulatory circuit driving the asymmetric-to-symmetric division switch (Figure 7E): First, SPCH initiates and sustains the fast and reiterative ACDs of a meristemoid. During the meristemoid-to-GMC transition, MUTE directly induces SMR4, which directly associates with CYCD3;1 (and likely with CYCD3;2) and inhibit of a CYCD3;1-CKDA;1 complex to terminate amplifying ACDs. At the same time, MUTE directly induces CYCD5;1. Because CYCD5;1 is not directly inhibited by SMR4, the final SCD can start even in the presence of SMR4. SMR4 may fine-tune the SCD by being able to inhibit the later-expressed CYCD7;1, which is likely complexed with CDKA;1. The endogenous expression of SMR4 disappears immediately after the execution of SCD (Figure 2F), hence the robust differentiation of stomata ensured. SMR8 has partially redundant role with SMR4 and is weakly induced by both SPCH and MUTE (Figure 1) as such, SMR8 is likely participating in part of this fine-tuning this transition. Unlike CycD3s and CycD7;1, CycD5;1 lacks part of the core domain (Strzalka et al., 2015), which may explain the differential SMR4 binding.

In addition to CKIs, Rb protein negatively regulates G1/S transition (Bertoli et al., 2013). The plant RETINOBLASTOMA RELATED (RBR) protein functions as key cell cycle regulators during stomatal development, and its reduced expression confers excessive proliferative ACDs within the stomatal cell lineages, in part due to dysregulated *SPCH* expression (Borghi et al., 2010; Weimer et al., 2012). Whereas both CYCD3;1 and CYCD7;1 possess LxCxE RBR-binding motif, CYCD5;1 bears a variant motif, which may compromise the RBR association (Vandepoele et al., 2002). Thus, CYCD5;1’s unique activity to execute the single SCD might involve the lack of negative regulation by RBR. Interestingly, RBR regulates the expression and activities of stomatal bHLH proteins, SPCH and FAMA, respectively (Matos et al., 2014; Weimer et al., 2012) but not MUTE. Thus, the commitment to differentiation by MUTE-orchestrated network may be inherently resilient to inhibition at G1/S transition.

Our study showed that extended G1-phase by stomatal-lineage overexpression of SMR4 conferred irregular-shaped meristemoids and eventual differentiation of stomata with skewed guard cells. Some guard cells exhibit a jigsaw-puzzled shape, which is indicative of pavement cell-like characteristics (Figures 4, 5). Thus, without timely execution of an ACD, the stomatal precursor cell can adopt hybrid identity of a guard cell and pavement cell. An intrinsic polarity protein BASL ensures that only one of the two daughter cells, the meristemoid, maintains high SPCH levels, thereby able to reiterate proliferative ACDs (Dong et al., 2009). The remaining daughter cell readily loses SPCH protein and differentiate into a pavement cell. This process involves a dynamic subcellular re-localization of BASL protein between the nucleus and polarly-localized cell cortex, the latter activates MAP kinase cascade that inhibits SPCH protein accumulation via phosphorylation (Zhang et al., 2016; Zhang et al., 2015). It is not known whether the cell cycle phase influences BASL behaviors, but our work implies that could be the case.

The SMR4-mediated cell cycle deceleration during the meristemoid-to-GMC transition mirrors the fundamental importance of G1-phase extension for cell fate decision and differentiation during development (Dalton, 2015; Liu et al., 2019). The direct role of MUTE to execute both termination of proliferative asymmetric divisions and orchestration of the single terminal symmetric division occurs through interwoven regulation of core cell cycle drivers and a braker. Understanding how cell cycle machineries in turn regulate the precise expression of MUTE, which likely involves epigenetic state changes, will provide a full picture of cell cycle control of cell fate specification in plants.

## ACKNOWLEDGEMENTS

We thank Bénédicte Desvoyes and Crisanto Gutierrez for *PlaCCI*, Lieven De Veylder for *proSMR1::nls-GFP*, Daisuke Kurihara for *proRPS5A::H2B-GFP*, and ABRC for GFP-Lti6B and *SMR8* T-DNA lines. This work is supported by JSPS KAKENHI Grant-in-Aid for Scientific Research on Innovative Areas (17H06476), WPI-ITbM operational funds, and the start-up funds from the UT Austin, Molecular Biosciences to KUT. KUT acknowledges the support from Howard Hughes Medical Institute and Johnson & Johnson Centennial Chair of Plant Cell Biology at the UT Austin. SKH was supported by the Young Leader Cultivation Program from the Institute of Advanced Research, Nagoya University.

## AUTHOR CONTRIBUTIONS

Conceptualization, S.-K.H. and K.U.T.; Experimental Design, S.-K.H, E.-D.K. and K.U.T.; Performance of experiments, S.-K.H., J.Y., M.A., R.I., T.S.; Bioinformatics Analysis, S.-K.H., S.K., E.-D.K. and K.U.T.; Visualization, S.-K.H., J.Y., K.U.T; Writing-Original Draft, S.-K.H. and K.U.T.; Writing-Review & Editing, S.-K.H., J.Y., E.-D.K., K.U.T; Project Administration, K.U.T.; Funding Acquisition, K.U.T.

## MATERIALS AND METHODS

### Plasmid Construction and Plant materials

For a detailed information of constructs generated in this study, see Table S4. Primers used for plasmid constructs were listed in Table S5-7. For generation of transgenic Arabidopsis, plasmid constructs were electroporated into Agrobacterium (GV3101/pMP90) and subsequently transformed by floral dipping. Loss-of-function mutant of *SMR4* was generated by CRISPR by using pKAMA-ITACHI Vector (Addgene: 85808) as described previously (Tsutsui and Higashiyama, 2017). Briefly, primers for sgRNA were designed by the CCTop - CRISPR/Cas9 target online predictor (Stemmer et al., 2015). Primers were annealed and inserted into pKI1.1R vector cut with *AarI*. Resulting construct was introduced into wild-type Col-0 plants. T1 plants were screened by Hygromycin resistance. Six T1 lines were selected and sequenced to check whether mutations were introduced. One of the two sgRNAs was successful for generating mutations. In T2 generation, seeds that do not show OLE1-RFP signal were selected to exclude plants harboring transgene in the genome. We established two independent homozygous lines that contain 1 bp deletion (*smr4-1*) or 1 bp insertion *(smr4-2)* at *SMR4* gene (Figure S4). Primers used single guide RNA (sgRNA) for *SMR4* were listed in Table S6. T-DNA insertion mutants of *SMR8, smr8-1* (SALK_126253), *smr8-2* (SALK_074523) were obtained from ABRC. Their genotype and transcript reduction were confirmed. Genotyping- and qRT-PCR primers were listed in Table S7. All plant materials used in this study were listed in Key Resource Table.

### Plant growth condition and estradiol treatment

Arabidopsis accession Columbia (Col-0) was used as wild-type. Sterilized seeds were grown on half strength of Murashige and Skoog (MS) media with 1% sucrose at 22°C under continuous light. 10∼14-day-old seedlings grown on MS media were transplanted to soil. For phenotyping of transgenic plants, either T1 or T2 plants were grown on ½ MS agar media containing hygromycin (20 μg/ml). For the complementation test of SMR4, T3 homozygous plants of *proSMR4::HA-SMR4* were germinated on ½ MS agar media with 1% sucrose and were imaged at 5-day post germination (dpg).

### Confocal microscopy

For confocal microscopy, cell peripheries of seedlings were visualized with either propidium iodide (Sigma, P4170) or FM4-64 (Invitrogen, T13320). Images were acquired using LSM800 (Zeiss) or SP5-WLL (Leica) using a 63x water lens. The Zeiss LSM800 was used to image the GFP and RFP reporter with excitation at 488 nm and an emission filter of 490 to 546 nm, and with excitation at 555 nm and 583-617 nm emission range, respectively. PlaCCI lines (Desvoyes et al., 2020) were imaged using SP5-WLL with the following conditions: CFP, excitation at 458 nm and emission from 468 to 600 nm; GFP, excitation at 488 nm and emission from 490 to 546 nm; YFP, excitation at 514 nm and emission from 524 to 650 nm; mCherry, excitation at 560 nm and emission from 590 to 650 nm. Signals were visualized sequentially using separate HyD detectors. DIC images were taken to delineate the cell outlines (shown in magenta). Raw data were collected with 1024 × 1024 pixel image and imported into Fiji-ImageJ to generate CYMK images using the channel merge function. The time-lapse were collected at 30-min intervals using a 20x lens (Peterson and Torii, 2012; Qi et al., 2017). Raw images were imported into Fiji-ImageJ to generate time projections using the Stacks function.

### Quantitative analysis of epidermal phenotype

For quantitative analysis of leaf epidermis of *smr* mutants, confocal images were taken at 4 days after germination. Preparation of images was done as described previously (Houbaert et al., 2018). For counting epidermal cell types, the number of stomata, number of stomatal lineage precursors (meristemoids and GMC), total epidermal cells (stomata, meristemoids, GMC and pavement cells) and stomata index (number of stomata / number of total epidermal cells x100) were calculated by counting cell types in an area of 0.45 mm^2^ (0.67 mm x 0.67 mm) at the developmental stage indicated with the cell counter plug-in in Fiji. The epidermal cell areas of *smr* mutants were color-coded depending on the area calculated using ROI_Color_Coder with a range of min-max (0-4000) in Fiji. The epidermal cells were subdivided into 9 groups according to their size. One representative image from each genotype was analyzed and the cell size distribution was then calculated from 499 cells for Col-0 plants, 659 cells for *smr4-1*, 662 cells for *smr4-2*, 601 cells form *smr8-1*, 611 cells form *smr8-1* and 755 cells for *smr4-1 smr8-1* double mutant. Guard cells were excluded for the cell size measurement.

For cell size and circularity measurement of stomatal lineage precursors, images of *proTMM::GUS-GFP* were set to grayscale and GFP expressing cells were colored in black while other cell area in white by photoshop, then the images were imported to Fiji. Imported images were subjected to Images > Threshold; Analyze > analyze particle. Shape descriptors box has to be checked in “Set measurement” under the Analyze tab to get circularity values from the selected cell area. For the meristemoid size in *mute* mutant background was measured by the same methods.

### cDNA preparation and qRT-PCR

For chemical treatment, plants were grown on media containing either 10uM β-estradiol (Sigma, E8875) or DMSO. For time-course induction, estradiol-inducible MUTE or SPCH seeds were sown on 1/2 MS media, and subjected to stratification at 4°C for two to three days then grown for the four to five days under continuous light. Subsequent steps were performed as described Han et al., 2018 (Han et al., 2018). RNA was isolated using RNeasey Plant Mini Kit (Qiagen, 74904). 0.5 μg of RNA was converted to cDNA using ReverTra Ace™ qPCR RT Master Mix with gDNA Remover (TOYOBO, FSQ-301) according to the instructions of the manufacturer. qRT-PCR was performed as described in Han et al., 2018 (Han et al., 2018) using KAPA SYBR® FAST qPCR Kit Master Mix on LightCycler® 96 instrument (Roche). Relative expression was calculated by dividing *ACTIN2* gene expression over the specific-gene expression and the fold change was calculated by dividing estradiol expression over DMSO (mock) expression at each time point indicated. See Table S7 for primer sequences used for qRT-PCR.

### Chromatin Immunoprecipitation Sequencing (ChIP-seq)

For MUTE ChIP-seq experiments, transgenic plants *proMUTE::MUTE-GFP scrm-D* were prepared as described previously (Han et al., 2018) with following modifications. To shear the DNA, Bioruptor (Diagnode) was used, 30 sec on and 30 sec off cycle 15∼18 times. Immunoprecipitation against GFP was performed using anti-GFP antibody (Abcam, ab290, Lot. GR318425-1). DNAs from the immuno-complex was purified by kit (Zymo Research, D5205). The half of the purified DNA was subjected to library preparation using ACCEL-NGS® 2S PLUS DNA LIBRARY KIT with 2S Set A MID Indexing Kit (Swift bioscience, 21024, 26148) for next generation sequencing. Quantitative PCR (qPCR) was carried out using gene specific primers (Table S7) to confirm the library construction. The qPCR was run using KAPA SYBR® FAST qPCR Kit Master Mix on LightCycler® 96 instrument (Roche) as described previously described (Han *et al*., 2018). Three biological replicates were used for MUTE ChIP-seq experiments. Size distribution of the libraries was validated by 2100 Bioanalyzer (Agilent). The prepared libraries (Col input and IP, MUTE-GFP input and IP with three replicates) were sequenced 35 bp paired-end in length with 30million coverage per sample on Illumina NextSeq 500 system. ChIP mapping and peak calling were performed as described in Feng et al., 2012 (Feng et al., 2012). Output reads were mapped to the TAIR10 genome assembly using bowtie2 and resulting bam files were sorted and indexed via samtool. The sorted bam files were subjected for MACS peak calling (version 2.1.0.20140616) (Table S2). Bedgraph file was generated and visualized in igv browser. Gene Ontology enrichment analysis was performed using GENE ONTOLOGY (http://geneontology.org/) combined with manual curation to remove redundant terms. Genes increased by MUTE more than Log2 FC 0.4 and targeted by MUTE were tested. Following multiple hypothesis testing correction (Bonferroni-correction), GO term with FDR <0.05 were called significantly enriched (Table S3). The ChIP-seq data generated in this study are deposited to the NCBI Gene Expression Omnibus (GEO) with an accession number GEO: GSE173338.

### Measurement of DNA content and nuclei size

1st true leaves were harvested from 16 day old plants and fixed in a solution of 9:1 (v/v) ethanol:acetic acid for overnight. For DAPI staining, tissues were rehydrated with ethanol series. DAPI (4’6-diamidino-2’-phenylindole) staining was done in 5µg/ml final concentration for 15 minutes in dark. Nuclear DAPI fluorescence was excited at 405nm captured with 410 - 470 nm emission range. DAPI stained nuclei area from the guard cells was selected and measured by using FIJI software. Wild-type and two independent T2 *proPOLAR::SMR4* transgenic lines were used. number of guard cells measured, 129 (WT) and 234 (*proPOLAR::SMR4*). 10 or 11-day-old cotyledons from *proRPS5A::H2B* (Maruyama et al., 2013) transgenic plants were also imaged to measure the nuclei size of guard- and pavement cells. The area of nuclei (GFP) was selected and measured from the z-stack projection images using FIJI software. Number of guard cells and pavement cells measured, 155 and 108 (wild type); 191 and 103 (*proPOLAR::SMR4 proRPS5A::H2B*).

### Yeast two hybrid assay

Y2H assays were performed using the Matchmaker™ Two-Hybrid System (Clontech). Bait (pGBKT7) and prey (pGADT7) constructs were co-transformed into the yeast strain AH109 according to manufactural instruction (Clontech). The resulting transformants were spotted on SD (−Leu, −Trp) and SD (−Trp, −Leu, −His) selection media containing different concentration of 3-amino-1,2,4-triazole (Sigma, A8056) as previously reported (Putarjunan et al., 2019). All constructs and primer information are listed in the Tables S4 and S5.

